# A New Interpretation for Oxygen Atom-Transfer Reactions for the Berg-Holm Oxo-Molybdenum Enzyme Model: Evidence for a Highly Active Oxygen Atom Transfer Acceptor

**DOI:** 10.1101/2025.01.08.631522

**Authors:** Jeremy M. Berg

**Affiliations:** Department of Computational and Systems Biology, University of Pittsburgh School of Medicine, Pittsburgh, PA 15213

## Abstract

In 1984, a synthetic model system for certain molybdenum oxotransferase enzymes was reported. These reports claimed that an oxygen atom could be extracted from a designed dioxomolybdenum(VI) complex to produce a monoxomolybdenum(IV) complex without the formation of an oxo-bridged molybdenum(V) binuclear species. The reduced product was shown to accept oxygen atoms from substrates such as dimethylsulfoxide with substrate saturation kinetics. Fifteen years later, it was demonstrated that the reduced product was, in fact, the oxo-bridged molybdenum(V) binuclear species. Here, it is shown that the kinetic data can be reinterpreted in terms of rate-limiting disproportionation of the oxo-bridged molybdenum(V) binuclear species to form a highly reactive monoxomolybdenum(IV) complex. The second order rate constant for oxygen atom transfer from dimethyl sulfoxide to this complex is more than 100,000 times higher than those reported for other monoxomolybdenum(IV) complexes. The five-coordinate molybdenum sites in the dioxomolybdenum(VI) and presumed monoxomolybdenum(IV) complexes are quite similar to those observed for eukaryotic nitrate reductase enzymes and this model system shows relatively rapid reduction of nitrate through a similar mechanistic scheme.

## Introduction

In 1984, Professor R.H. Holm and I reported a synthetic reactivity model for the active sites of molybdenum-containing oxotransferases^1^. The key component of this system is MoO_2_(L-NS_2_) where L-NS_2_ is the doubly deprotonated form of 2,6-Bis(2,2-diphenyl-2-mercaptoethyl)pyridine. This crystal structure of this compound^2^ revealed a trigonal bipyramidal structure (Figure 1).

**Figure 1.**
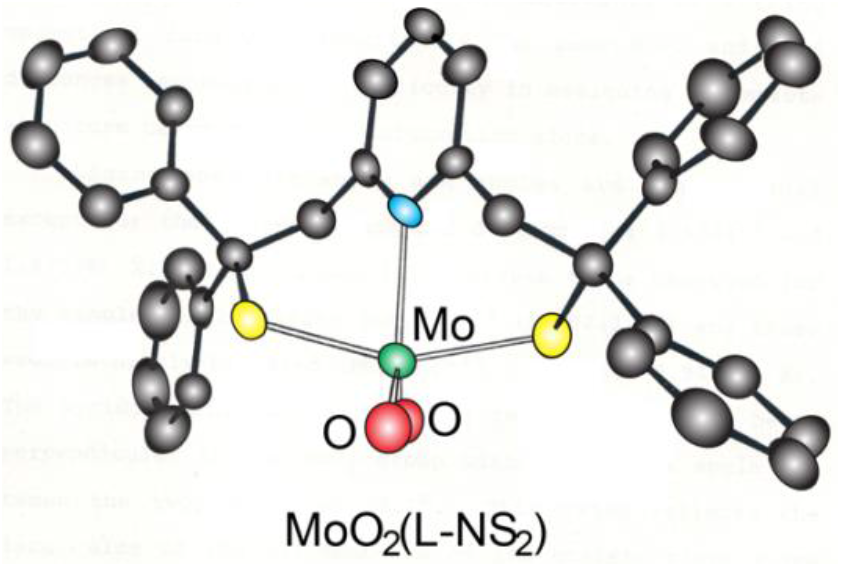
The structure of [2,6-Bis(2,2-diphenyl-2-thioethyl)pyridinato]dioxomolybdenum(VI), (MoO_2_(L-NS_2_) (redrawn from reference 2).

Reaction of solutions of orange MoO_2_(L-NS_2_) with Ph_3_P in N-dimethylformamide (DMF) solution produced a purple product with clean isosbestic points in visible absorption spectra suggesting conversion to a single colored product^3^. This product, hereafter referred to as [MoO_2_(L-NS_2_)]_reduced_, could be isolated as a purple, microcrystalline product and OPPh_3_ was confirmed as the other product by ^31^P NMR spectroscopy^3^, supporting equation 1.

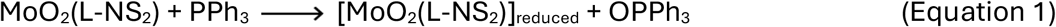

MoO_2_(L-NS_2_)]_reduced_ reacted with sulfoxides such as dimethylsulfoxide (Me_2_SO) to form MoO_2_(L-NS_2_) and Me_2_S^3^. Examination of the rate of this reaction as a function of Me_2_SO concentration revealed substrate saturation kinetics (Figure 2)^3^.

**Figure 2.**
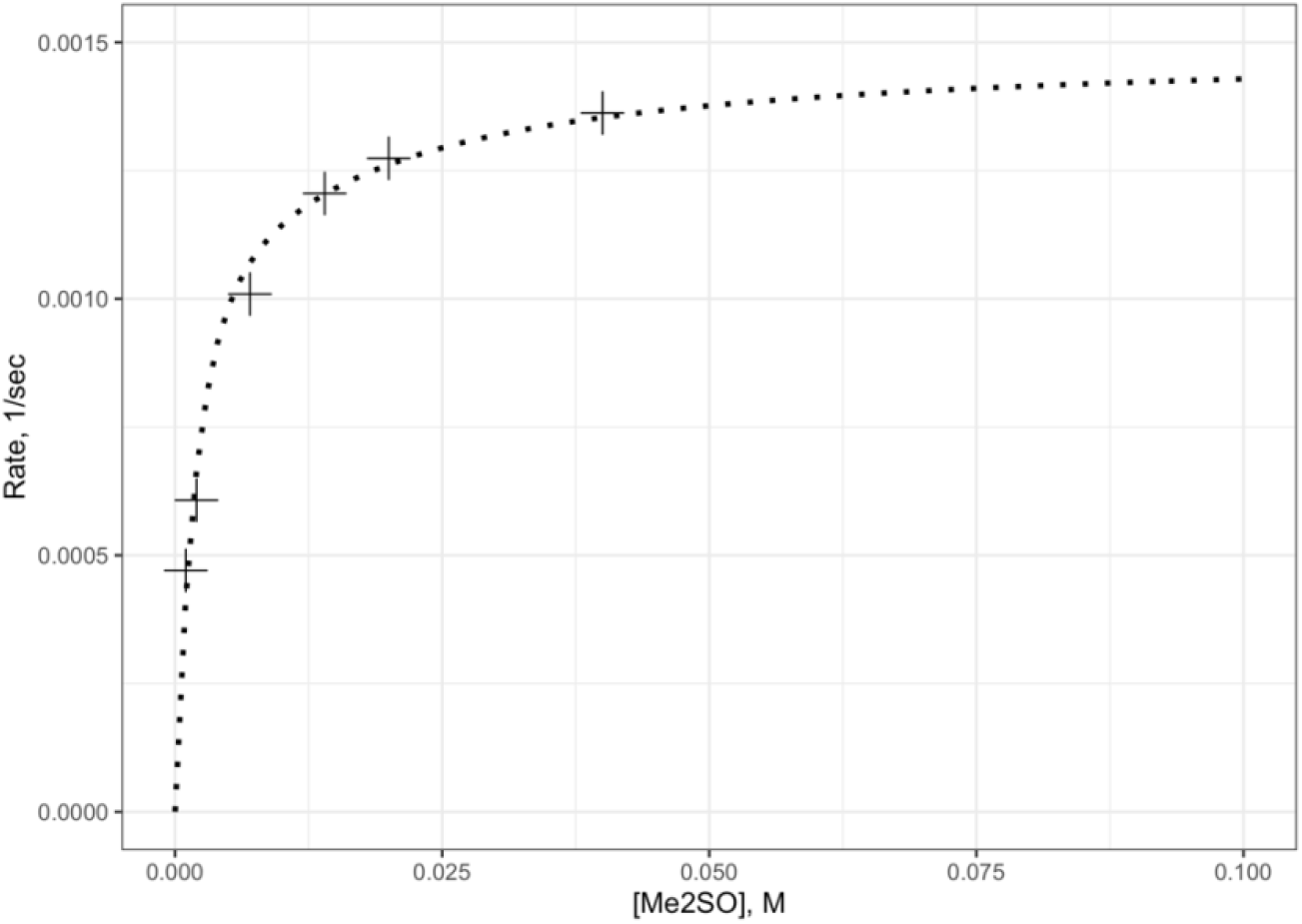
The kinetics of Me_2_SO reduction by [MoO_2_(L-NS_2_)]_reduced_ as a function of Me_2_SO concentration showing substrate saturation with a maximal rate of approximately 0.0015 s-1 (redrawn from reference 3).

Based on elemental analysis and this reactivity, the structure of [MoO_2_(L-NS_2_)]_reduced_ was assigned as MoO(L-NS_2_)(DMF)^2^. The alternative assignment as Mo_2_O_3_(L-NS_2_)_2_.2DMF, an oxo-bridged binuclear species was considered, but we deemed less likely based on the reactivity with Me_2_SO, the role of the sterically bulky ligand that had been designed in inhibit the formation of such oligomeric species, and other unpublished evidence. Despite considerable effort, crystals suitable for structure determination could not be obtained.

Based on these observations, we proposed a reaction scheme^1^ for the oxygen atom-transfer reactions (Figure 3).

**Figure 3.**
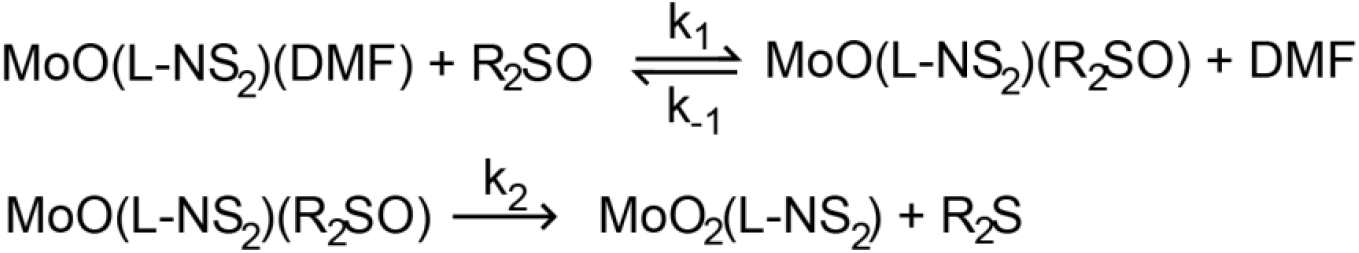
The initially proposed reaction scheme for sulfoxide reduction by [MoO_2_(L-NS_2_)]_reduced_ involves sulfoxide binding to the molybdenum(IV) center, followed by relatively slow oxygen atom transfer to form MoO_2_(L-NS_2_) and the corresponding sulfide.

The oxygen atom-transfer reactions could be performed catalytically more than 100 cycles^3^.

Reactions with a range of sulfoxide substrates including biologically relevant substrates such as methionine sulfoxides and d-biotin-d-sulfoxide were also examined with similar results^3^. All showed 1.36 – 1.70 × 10^−3^ s^-1^ limiting rates but somewhat more variable substrate concentrations at half-maximal rates of 0.8 to 3.1 mM. These half-maximal concentrations values were interpreted in terms of equilibrium constants for displacement of DMF by substrate.

Another biologically relevant substrate NO_3_^-^ was also found to be reduced, but the reaction was not clean due to destruction of the molybdenum complex by the NO_2_^-^ product^3^. Subsequent experiments with a nitrite scavenger^4^ revealed a clean reaction with saturation kinetics with a limiting rate of 1.49 × 10^−3^ s^-1^ and a half maximal rate a nitrate concentration of near 1 mM.

Subsequently, additional substrates including (p-FC_6_H_4_)_2_SO and 3-fluoropyridine N-oxide were also examined^5^. Again, substrate saturation kinetics was observed with similar limiting rates of 1.4 to 1.6 × 10^−3^ s^-1^ but with substrate concentrations at half maximal rates differing by a factor of approximately 8.

Based on the relative independence of the apparent rate of substrate reduction on the nature of the substrate, and the lack of definitive structural characterization of the reduced product ([MoO_2_(L-NS_2_)]_reduced_), Holm and coworkers^6^ developed “an improved analogue system for the molybdenum oxotransferases”. This system was based on a very sterically hindered pyridine thiolate ligand (hereafter designated tBuL-NS) that formed a dioxomolybdenum(VI) complex Mo(VI)O_2_(tBuL-NS)_2_ that could be reduced to Mo(IV)O(tBuL-NS)_2_. Both the oxidized and reduced forms could be characterized crystallography. Mo(IV)O(tBuL-NS)_2_ could accept oxygen atoms from a variety of substrates. The reduction of Me_2_SO proceeded with a second-order rate constant of 1 × 10^−4^ M^-1^s-^1^.

Fifteen years after the original publications, Doonen et al.^7^ published “New Insights into the Berg-Holm Oxomolybdoenzyme Model”. The primary new insight involved the structural characterization of ([MoO_2_(L-NS_2_)]_reduced_, demonstrating that this product was not the proposed Mo(IV)O(L-NS_2_)(DMF) but rather was the DMF solvate of the binuclear complex Mo(V)_2_O_3_(L-NS_2_)_2_. The crystal structure of this complex demonstrated that ligand reorientation allowed the formation of a μ-oxo-binuclear species without steric interference of the gem-diphenyl groups^7^. The definitive characterization of the reduced product from the Berg-Holm oxotransferase as Mo(V)_2_O_3_(L-NS_2_)_2_ requires a reinterpretation of the reaction schemes within this system. In particular, the rate-limiting step for the reduction of sulfoxide and other oxygen atom donors is assigned not to the oxygen atom transfer reaction itself, but rather to the disproportion of Mo(V)_2_O_3_(L-NS_2_)_2_ to form Mo(VI)O_2_(L-NS_2_) and a molybdenum(IV) species, presumably Mo(IV)O(L-NS_2_)(DMF).

## Methods

Rate data for substrate reduction reactions for Me_2_SO^3^, Ph_2_SO^3^, d-biotin-d-sulfoxide^3^, NO_3_^-4^, (p-FC_6_H_4_)_2_SO^5^, and 3-fluoropyridine N-oxide^5^ were digitized from published figures. Rate data for substrate reduction reactions for carbobenzyloxy derivatives of L-methionine-d-sulfoxide and L-methionine-l-sulfoxide^8^ were digitized from reference 8. Rate data were adjusted by 8% or less so that the maximum rate was 0.0015 s^-1^. These corrections may reflect minor uncertainties in the concentrations of the molybdenum complex used.

Kinetic rate equations were analyzed with RStudio^9^ using the package deSolve^10^. Curve fitting was performed using chi square as a goodness-of-fit parameter. 95% confidence intervals were estimated from the covariance matrix derived from the fit.

## Results

The new proposed reaction scheme is shown in Figure 4.

**Figure 4.**
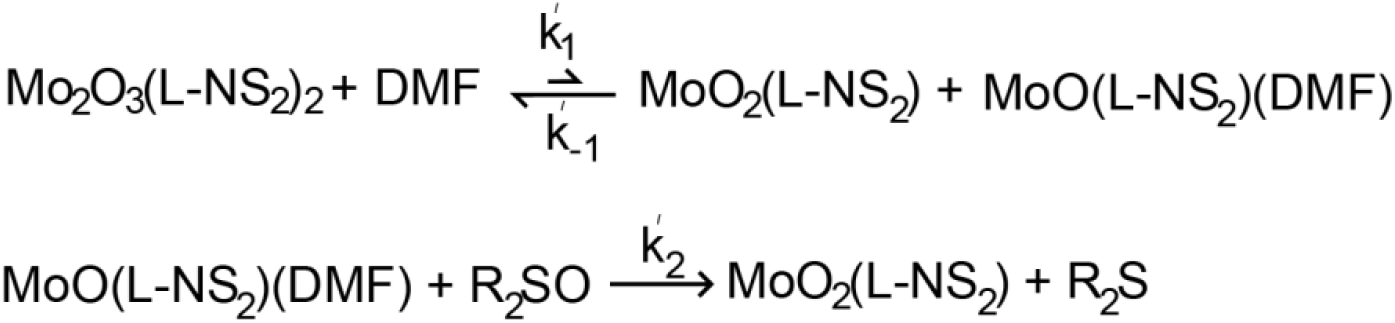
An alternative mechanistic scheme for sulfoxide reduction by ([MoO_2_(L-NS_2_)]_reduced_. The reaction begins with the binuclear Mo(V) complex which disproportionates slowly to produce the dioxomolybdenum(VI) complex and a reactive monoxomolybdenum(IV) complex that can accept an oxygen atom to reduce a sulfoxide to the corresponding sulfide.

According to this scheme, the rate of substrate reduction is limited by the rate of disproportionation. Substrate saturation kinetics is expected due to competition for the reactive monoxomolybdenum(IV) species between substrate and the dioxomolybdenum(VI) complex that accumulates over the course of the reaction.

The kinetic parameters can be estimated from several considerations. The disproportionation rate constant, k_1_’, must be approximately the observed limiting rate of 0.0015 s^-1^. A minimum value for the reverse rate constant (for the formation of Mo(V)_2_O_3_(L-NS_2_)_2_) can be estimated by the fact that the reaction does not proceed beyond this product to any appreciate event and clean isosbestic points are observed throughout the reaction. Simulation studies reveal that this requires k_-1_ to be greater than 100 M^-1^s-^1^. This estimate can be refined by fitting the observed reaction rate curves. The results of such fits are shown in Figure 5 for the reduction of Me_2_SO.

**Figure 5.**
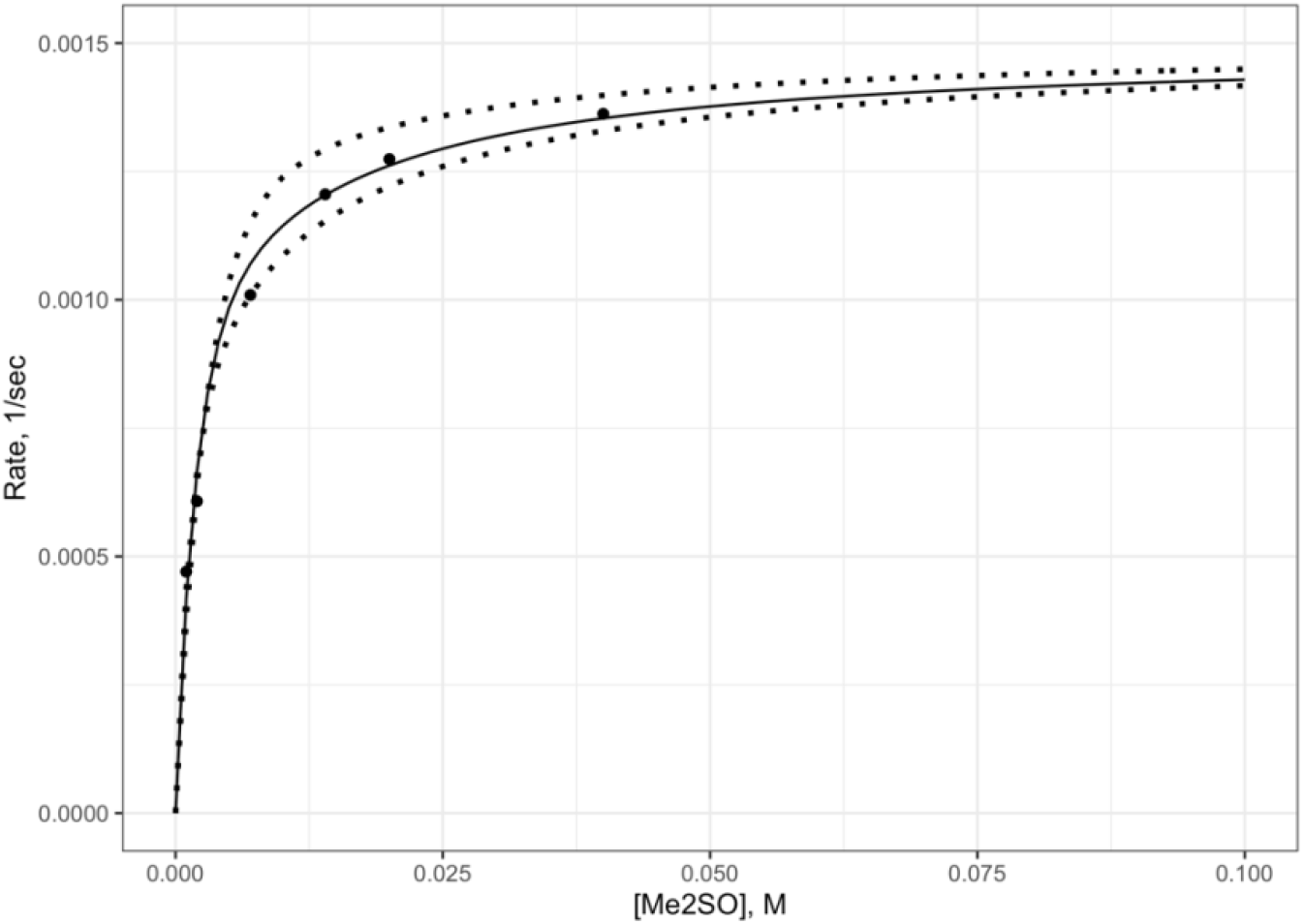
The rate of dimethyl sulfoxide reduction by Mo_2_O_3_(L-NS_2_)_2_ fit with k_1_’ = 0.0016 s^-1^, k_-1_’ = 4000 M^-1^ s^-1^, and k_2_’ = 11 M^-1^ s^-1^. For comparison, fits with k_-1_’ = 2000 M^-1^ s^-1^ and k_-1_’ = 6000 M^-1^ s^-1^ with other parameters optimized, are shown (dotted lines) to demonstrate the sensitivity of the curves to the k_-1_’ parameter.

Substrate reduction data are available for 8 substrates. These data were fit using a global model with single values for k_1_’ and k_-1_’ and individual values of the oxygen atom transfer rate, k_2_’, for each substrate. The results are shown in Figure 6.

**Figure 6.**
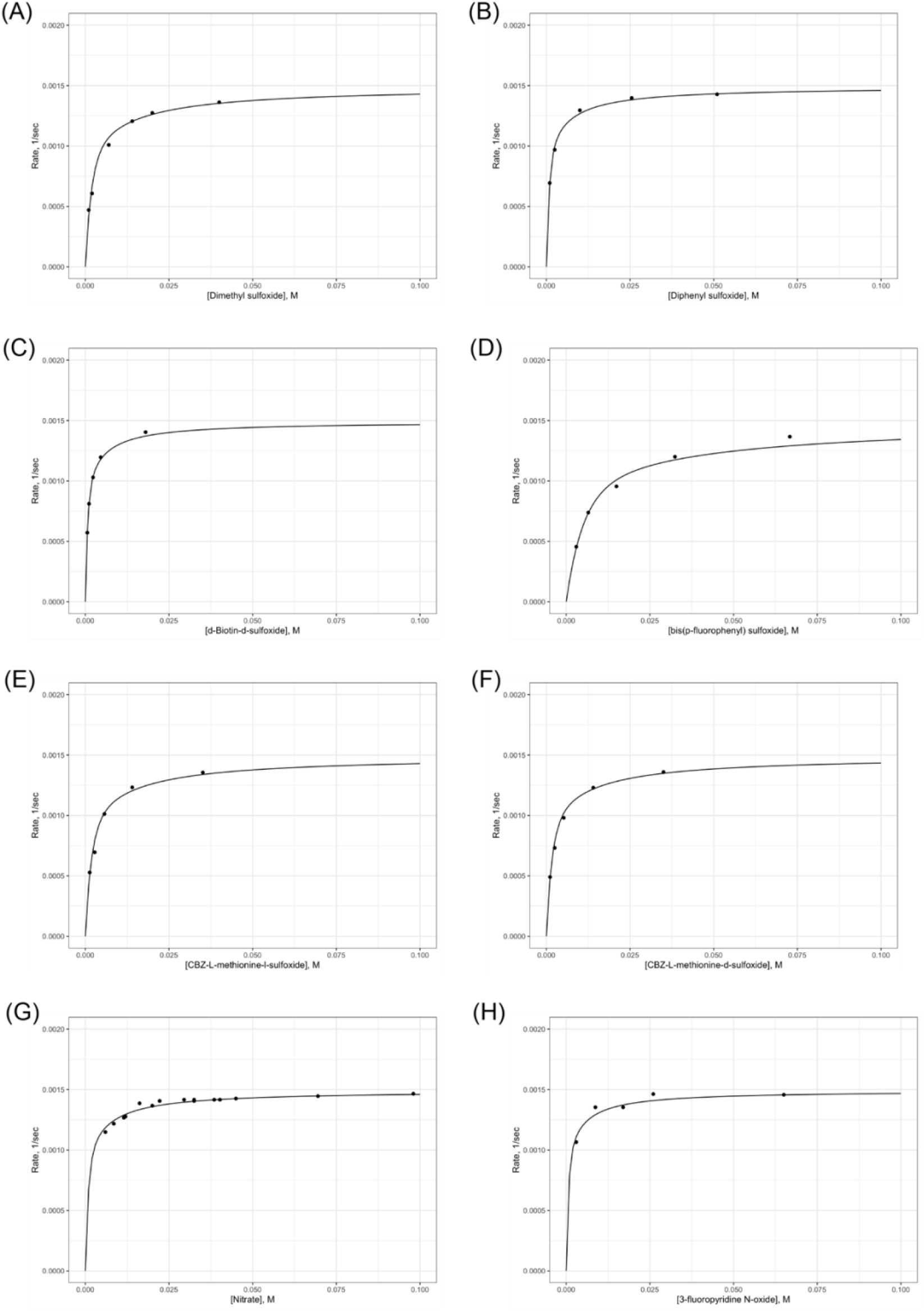
Observed versus calculated substrate reduction curves for eight substrates fit with single parameters for the disproportion reaction (k_1_’ = 0.0016 s^-1^ and k_-1_’ = 4000 M^-1^ s^-1^) with different rate constants for the oxygen atom transfer reaction for each substrate. (A) Me2SO; (B) Ph2SO; (C) d-Biotin-d-sulfoxide; (D) (p-FC_6_H_4_)_2_SO; (E) L-methionine-l-sulfoxide; (F) L-methionine-d-sulfoxide; (G) NO_3_^-^; (H) 3-fluoropyridine-N-oxide.

The results of this global fit are shown in Table 1.

**Table 1.** Rate constants from global fit of substrate reduction rate data.

| Rate constant | Substrate | Value | 95% Confidence interval |
| --- | --- | --- | --- |
| $k_1'$ | All | $0.0016 \text{ s}^{-1}$ | 0.00155 to 0.00165 |
| $k_{-1}'$ | All | $4000 \text{ M}^{-1}\text{s}^{-1}$ | 2600 to 5800 |
| $k_2'$ | Dimethyl sulfoxide | $11 \text{ M}^{-1}\text{s}^{-1}$ | 7 to 16 |
| | Diphenyl sulfoxide | $23 \text{ M}^{-1}\text{s}^{-1}$ | 20 to 27 |
| | d-Biotin-d-sulfoxide | $29 \text{ M}^{-1}\text{s}^{-1}$ | 26 to 33 |
| | Bis(p-fluorophenyl) sulfoxide | $4 \text{ M}^{-1}\text{s}^{-1}$ | 1 to 7 |
| | CBZ-L-methionine-l-sulfoxide | $11 \text{ M}^{-1}\text{s}^{-1}$ | 7 to 14 |
| | CBZ-L-methionine-d-sulfoxide | $12 \text{ M}^{-1}\text{s}^{-1}$ | 8 to 14 |
| | Nitrate | $23 \text{ M}^{-1}\text{s}^{-1}$ | 20 to 26 |
| | 3-fluoropyridine-N-oxide | $31 \text{ M}^{-1}\text{s}^{-1}$ | 23 to 39 |

## Discussion

With the definitive identification of the product of reduction of Mo(VI)O_2_(L-NS_2_) as Mo(V)_2_O_3_(L-NS_2_)_2_, the substrate saturation kinetics behavior observed for this system was reinterpreted in terms of rate dissociation of Mo(V)_2_O_3_(L-NS_2_)_2_ to form the reactive species Mo(IV)O(L-NS_2_)(DMF). Using this kinetic scheme, rate data for eight substrates could be fit well, yielding the rate constant for the combination of Mo(VI)O_2_(L-NS_2_) and Mo(IV)O(L-NS_2_)(DMF) and rate constants for the oxygen atom transfer reaction for each substrate.

The rate constant for the combination of Mo(VI)O_2_(L-NS_2_) and Mo(IV)O(L-NS_2_)(DMF) of 4000 M^-1^s^-1^ is comparable to the value of 2350 M^-1^s^-1^, observed^11^ for the well-studied oxomolybdenum system Mo(VI)O_2_(S_2_CNEt)_2_/ Mo(V)_2_O_3_(S_2_CNEt)_4_/ Mo(IV)O(S_2_CNEt)_2_^12^. The disproportionation rate constant of 0.0016 s^-1^ for Mo(V)_2_O_3_(L-NS_2_)_2_ is 3000-fold smaller than that for Mo(V)_2_O_3_(S_2_CNEt) _4_^11^.

The oxygen atom transfer rate constant is lowest for Bis(p-fluorophenyl) sulfoxide with its electron-withdrawing substituents. The oxygen atom transfer rates constants are essentially identical for Me_2_SO and the two isomers of CBZ-L-methionine sulfoxide, consistent with the very similar electronic and steric environments for these sulfoxides. The largest rate constant is observed for the pyridine-N-oxide substrate that had a weaker X-O bond^13^.

The kinetics of oxygen atom transfer had been reported^13^ for the MoO_1-2_(tBuL-NS)_2_ system for three of the substrates analyzed herein: Me_2_SO, Ph_2_SO, and 3-fluoropyridine-N-oxide with rate constants of 1.0 × 10^−4^ M^-1^s^-1^, 3.1 × 10^−4^ M^-1^s^-1^, and 6.5 × 10^−2^ M^-1^s^-1^, respectively. The ranked order of these rate constants are the same as those observed for MoO(L-NS_2_)(DMF). However, the rates for MoO(tBuL-NS)_2_ are substantially lower than those deduced for MoO(L-NS_2_)(DMF) with ratios of 110000, 74000, and 480. The most obvious explanation for these large differences in rates is the fact that MoO(tBuL-NS)_2_ is five coordinate with no open coordination site whereas the reduced form of MoO_2_(L-NS_2_) is likely MoO(L-NS_2_)(DMF) with a bound solvent displaceable by substrate. This difference may also account for the less steep dependence of the rate on the X-O bond strength^13^. The rate constant for Me_2_SO reduction^12^ by MoO(S2CNEt_2_)_2_ of 0.00016 M^-1^s^-1^ is 69000-fold smaller than that for MoO(L-NS_2_)(DMF), also supporting the benefits of an open coordination site on the monoxomolybdenum(IV) species.

The MoO_2_(L-NS_2_) system was designed and characterized prior to the knowledge of the three-dimensional structure of any molybdenum-containing enzyme. Information about the molybdenum coordination environments was derived from X-ray absorption spectroscopy. However, many such enzyme structures are now known^14^. The structures for various dimethyl sulfoxide and methionine sulfoxide reductases show active sites with molybdenum centers coordinated to four thiolates from two molybdopterin cofactors with only a single oxo group in the oxidized form. This difference in structure makes the present model system less relevant to these enzymes. In contrast, the structure of eukaryotic nitrate reductase^15^ shows a molybdenum site that is remarkably similar to that in this model system. In the oxidized form of nitrate reductase, the molybdenum is coordinated two thiolates from molybdopterin, one thiolate from a cysteine residue, and two oxo groups in an approximately trigonal bipyramidal structure with one thiolate from molybdopterin and the cysteine thiolate in axial positions. This is quite similar to the molybdenum site in MoO_2_(L-NS_2_) with its trigonal bipyramidal structure with a pyridine nitrogen in place of one of the molybdopterin thiolates. Moreover, the reaction scheme proposed for this enzyme is quite like that proposed for the model system (neglecting the formation of the oxo-bridged binuclear species) as shown in Figure 7. The occurrence of five-coordinate rather six-coordinate species throughout the reaction cycle may be responsible for the relatively rapid rates compared with other model systems.

**Figure 7.**
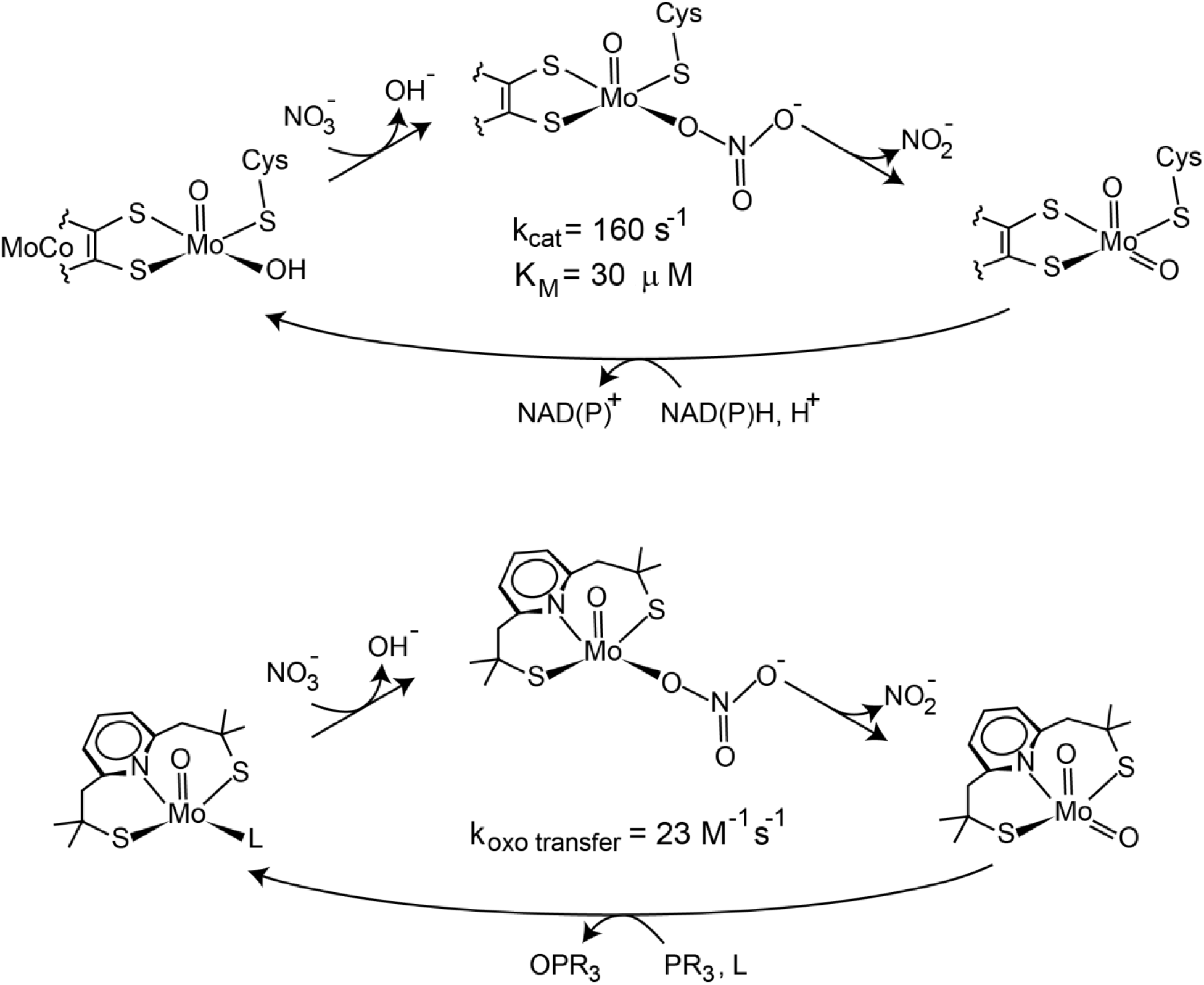
A comparison of the reaction scheme for eukaryotic nitrate reductase (adapted from reference 15) and that deduced for MoO_1,2_(L-NS_2_).

This kinetic scheme suggests that if MoO_2_(L-NS_2_) could be converted completely to MoO(L-NS_2_)(DMF), the reduced species would be a quite effective oxygen atom acceptor. For other systems that form oxo-bridged molybdenum(V) binuclear species reversible, it has been possible to drive the reactions through this intermediate for form monoxomoybdenum(IV) species. For example, treatment of MoO_2_(S_2_CNEt)_2_ withPh_3_P or other phosphines results in an initial buildup of Mo_2_O_3_(S_2_CNEt)_4_, followed by consumption of this to form MoO(S_2_CNEt)_2_. However, the rate constant for the disproportionation of this oxo-bridged binuclear species is 4.9 s^-1^, more than 3000 times than that for Mo(V)_2_O_3_(L-NS_2_)_2_.

If MoO(L-NS_2_)(DMF) could be prepared in pure form, it would be expected to be a very active oxygen atom acceptor. Of course, the MoO_2_(L-NS_2_) product formed would combine with the the MoO(L-NS_2_)(DMF) starting material to form Mo_2_O_3_(L-NS_2_)_2_. The expected accumulation of Mo_2_O_3_(L-NS_2_)_2_ over time for the reaction of 0.1 mM MoO(L-NS_2_)(DMF) with 1 equivalent of Me_2_SO is shown in Figure 8.

**Figure 8.**
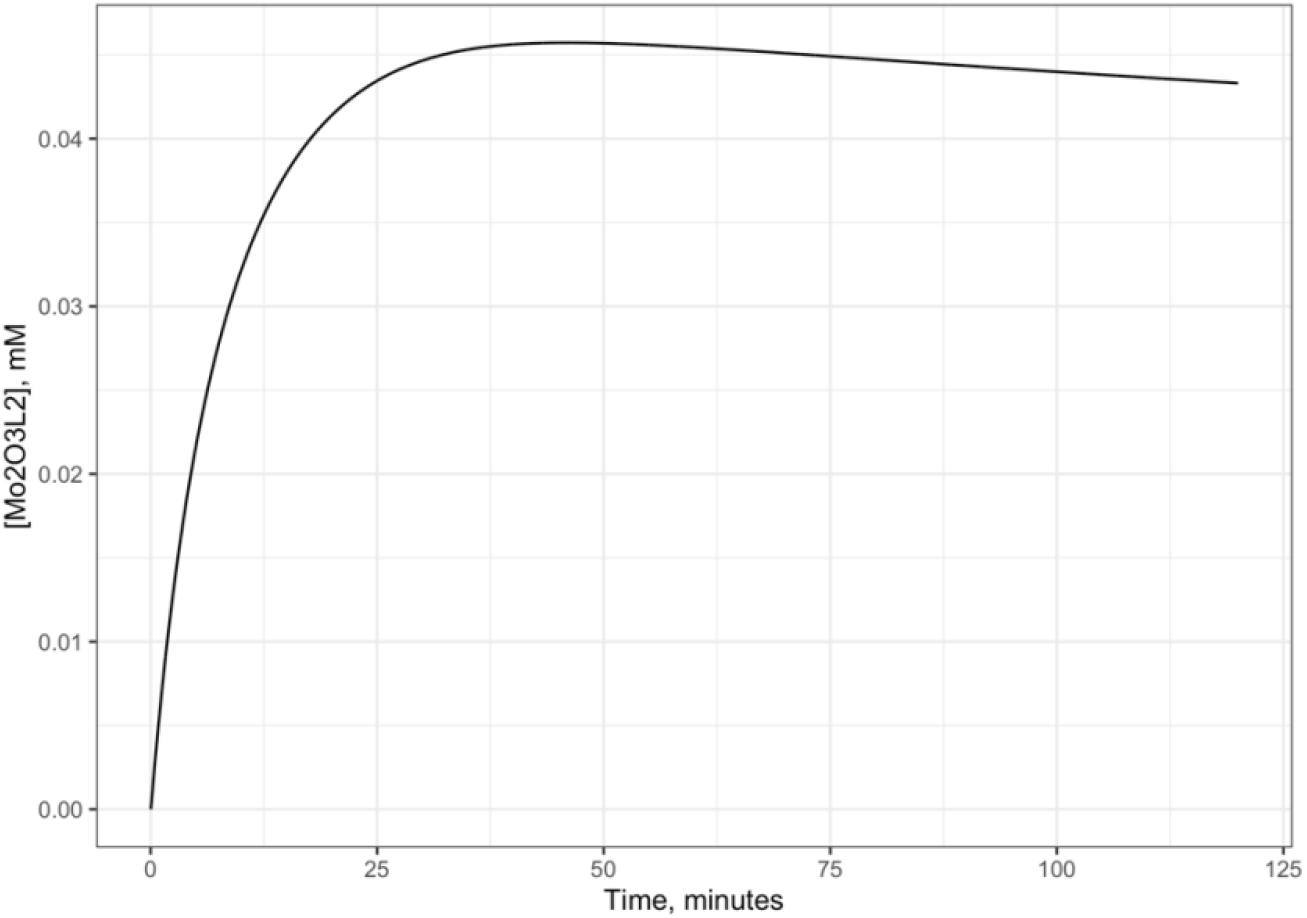
The predicted time course for treatment of 0.1 mM MoO(L-NS_2_)(DMF) with one equivalent of Me_2_SO at room temperature. The oxo-bridged molybdenum binuclear species accumulates rapidly over 20 minutes until most of the MoO(L-NS_2_)(DMF) has combined with the MoO_2_(L-NS_2_) formed. The binuclear species is then slowly consumed in a process limited by the rate of disproportionation.

The preparation of Mo(L-NS_2_)(DMF) has been attempted using higher temperatures and longer reaction times, revealed unanticipated reactivity of the L-NS_2_ ligand in the context of the Mo_2_O_3_((L-NS_2_) binuclear complex^16^. Alternative synthetic approaches beginning with Mo(IV) starting materials might be more fruitful.

## Conclusions

The kinetics of substrate reduction reactions by the reduced form of the Berg-Holm oxomolybdoenzyme model has been reinterpreted in light of the proof that this reduced species is Mo(V)_2_O_3_(L-NS_2_)_2_. Excellent fits of the kinetic data were obtained using a simple model. The deduced rates of oxygen atom transfer reactions to Mo(IV)(L-NS2)(DMF) formed by disproportionation of Mo(V)_2_O_3_(L-NS_2_)_2_ are many orders of magnitude higher than those observed for other model systems. The five-coordinate molybdenum site structures and reaction scheme for this model system closely mimic those for eukaryotic nitrate reductase.

## Acknowledgements

This paper is dedicated to the memories of Professors Richard H. Holm and Jeremy R. Knowles.

